# Repeated sleep deprivation selectively reactivates hippocampal CA1 pyramidal neuron

**DOI:** 10.1101/2025.07.30.665413

**Authors:** Yutong Wang, Emily N. Walsh, Junko Kasuya, Colton Remedies, Jon Resch, Lisa C. Lyons, Ted Abel

**Author notes:** **Address for Correspondence:** Ted Abel, Ph.D., Department of Neuroscience and Pharmacology, Iowa Neuroscience Institute, Carver College of Medicine, University of Iowa, 169 Newton Road, 2314 Pappajohn Biomedical Discovery Building, Iowa City, IA 52242, United States. **Author Email Addresses:** Yutong Wang, Emily N. Walsh, Junko Kasuya, Colton Remedies, Jon Resch, Lisa C. Lyons, Ted Abel.

## Abstract

Sleep supports a variety of physiological processes, ranging from metabolic to immune system homeostasis, and plays a critical role in cognition and memory. A brief period of sleep loss impairs memory, particularly hippocampus-dependent memories, and alters molecular signaling and synaptic plasticity in the hippocampus. Studies have shown that sleep deprivation (SD), alters neuronal activation as indicated by broad changes in gene expression signatures and by the altered expression of c-Fos, an immediate early gene that functions as a molecular marker of neuronal activity. In the present study, we examined hippocampal subregion-specific c-Fos induction patterns via immunohistochemical staining. We find that CA1 pyramidal neurons exhibit the most robust c-Fos induction after SD. Using an activity-driven ribosomal tagging system and a repeated SD model, we labeled sleep deprivation activated CA1 neurons and observed a population of excitatory neurons in area CA1 that are reactivated by repeated SD. Using the c-Fos-RiboTag system that enables the isolation of ribosomes attached mRNA from labeled neurons, we performed fosTRAP-seq and identified activity-dependent gene expression changes in c-Fos+ CA1 neurons. Our results revealed that synapse organization, protein dephosphorylation, cellular response to endogenous stimulus (such as insulin) are upregulated, whereas mRNA processing and splicing being downregulated. In summary, our study provides a detailed view of the activation of hippocampal neurons after SD, revealing a subset of CA1 pyramidal neurons that have higher sensitivity to the effect of sleep loss, shown as reactivation during repeated SD, allows investigation of molecular changes in neurons specifically impacted by repeated sleep loss. Our work uncovers a population of CA1 pyramidal neurons that are sensitive to repeated sleep loss and sheds light on a possible connection between acute and chronic sleep loss at the cellular and molecular levels.

**Graphical Abstract:** 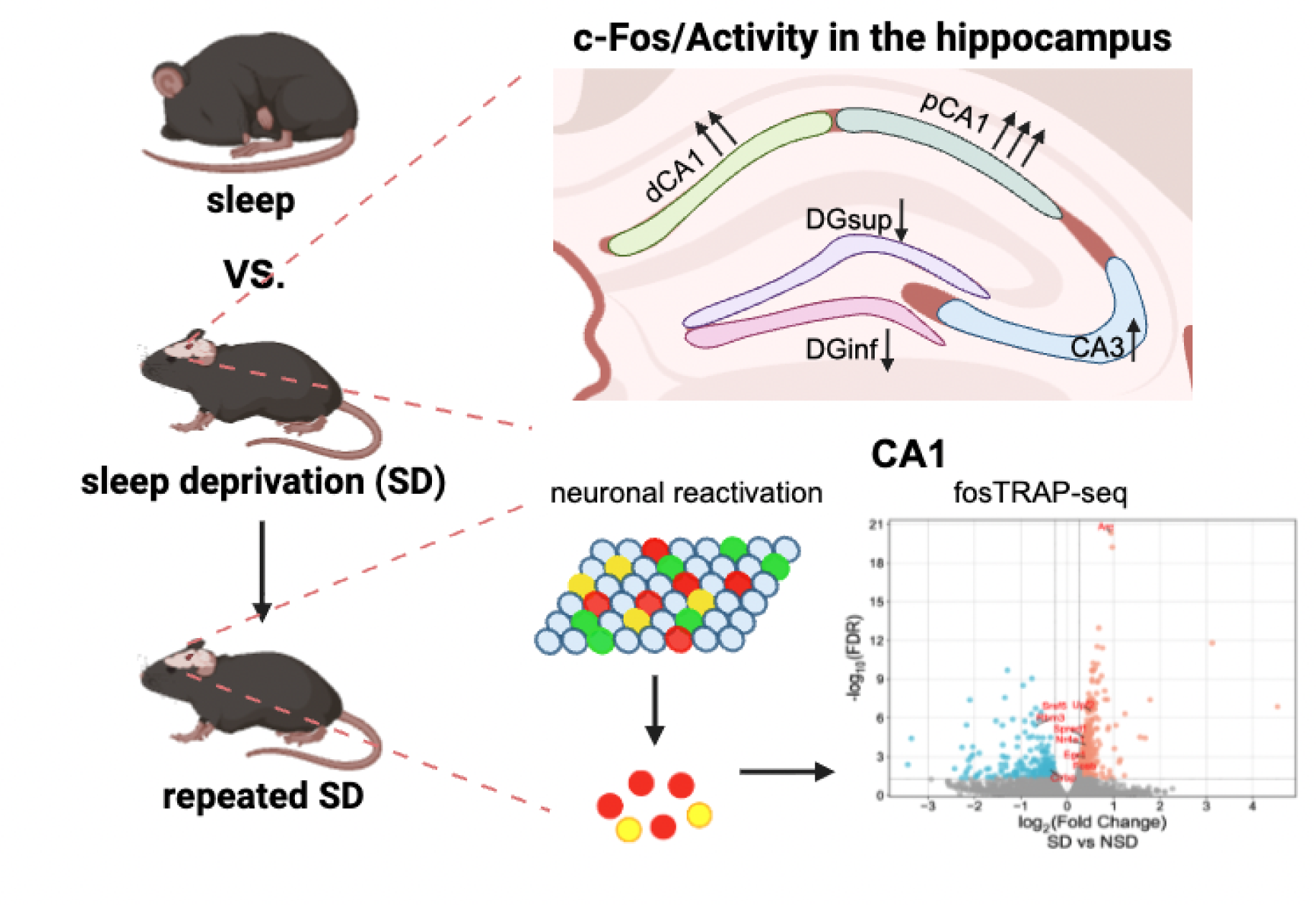

## INTRODUCTION

Sleep facilitates numerous physiological processes including cognition, learning, and memory consolidation (1). Over 30% of adults in the United States routinely experience insufficient sleep, leading to subsequent health problems (2, 3). Sleep loss has pronounced impacts on cognitive function, with hippocampus-dependent memories being especially vulnerable (4). The hippocampus plays a critical role in consolidating spatial, episodic, and contextual memories in rodents (5) and declarative memory in humans (6). It is comprised of subfields with specialized roles in memory: the Dentate Gyrus (DG) enables pattern separation (7), Cornu Ammonis 3 (CA3) integrates separation and completion (8), and Cornu Ammonis 1 (CA1) recodes CA3 outputs while linking to the neocortex for memory retrieval (9). Brief periods of sleep deprivation (SD) impact hippocampal function via impairments in the cAMP signaling and related long-lasting forms of synaptic plasticity (10–12), suppressed protein synthesis (13), and altered synaptic connectivity (14–16).

In humans, sleep loss can often span several nights or even weeks. Chronic insomnia is related to accelerated cognitive decline and is associated with increased risk of dementia in humans (17, 18). Work using rodents to model chronic sleep restriction (CSR) has demonstrated impairment of hippocampal synaptic plasticity and severe spatial memory deficits that are resistant to recovery sleep (19, 20). CSR also promotes tau hyperphosphorylation and amyloid-β accumulation in rodent models of Alzheimer’s Disease (21, 22). However, how repeated bouts of sleep loss affect neuronal activity in the hippocampus and how acute SD-induced impairments progress to CSR-promoted neurodegeneration remains not known.

A widely recognized marker of neuronal activity is the immediate early gene (IEG) c-Fos. The c-Fos protein is rapidly activated during synaptic plasticity associated with learning and memory (23), and its downstream effector genes, such as *Timp1* and *Mmp9*, regulate the neuronal structural and functional plasticity required for memory formation (23). c-Fos expression has been used as a marker to locate and identify the neuronal populations activated by sleep and extended wakefulness (24–27). While sleep is typically linked to low c-Fos expression and neuronal activity across most brain regions (24, 25), several hours of SD significantly increases c-Fos expression, particularly in the cortex, medial preoptic area, posterior hypothalamus, and the hippocampus (24–27). This brain-wide activation resembles the increased c-Fos expression pattern during spontaneous wakefulness. Given the anatomical and functional complexity of the hippocampus, this study aims to precisely map c-Fos expression patterns within hippocampal subregions after single and repeated SD exposures, and identify activity-driven gene expression changes within SD-activated neurons.

Our results suggest that neuronal activation, as measured by c-Fos expression, varies across the hippocampus following both single and repeated SD. Specifically, CA1 subregion showed the most pronounced increase in activation, while DG exhibited a decrease after both single and repeated SD. Applying the Target Recombination in Active Populations (fosTRAP) method in conjunction with the ribosome tagging strategy (RiboTag) (28, 29), we devised an active neuron labeling system that was amenable to analysis of individual gene expression. Through this c-Fos-RiboTag approach, we confirmed that a substantial proportion of CA1 pyramidal neurons were reactivated by repeated bouts of SD. As prior genomic studies identified translation-level changes in brain regions exhibiting increased c-Fos expression after SD, including the hippocampus (30–33), we further combining the c-Fos-RiboTag labeling system with Translating Ribosome Affinity Purification sequencing (TRAPseq) technique and mapped activity-dependent translatome changes in c-Fos+ CA1 neurons.

## METHODS & MATERIALS

### Animals

All experimental animals were 3- to 5-month-old male C57BL/6J mice obtained from the Jackson Laboratories (Catalog# 000664, Bar Harbor, ME). Mice were group housed (5 per cage) prior to the start of the experiment in standard ventilated cages with access to food (NIH-31 irradiated modified mouse diet #7913) and water *ad libitum.* Animals were maintained in the animal care facility at the University of Iowa on a 12h light/dark cycle, in a temperature- and humidity-controlled environment (21-22°C and 60-70%, respectively). All experiments were conducted in accordance with the standards established by the US National Institutes of Health Guidelines for Animal Care and use was approved by the Institutional Animal Care and Use Committee (IACUC) at the University of Iowa.

### Sleep deprivation

Total SD was performed acutely, for 5h beginning at Zeitgeber time (ZT) 0 using the gentle handling method, which involved lightly tapping and shaking the cage to keep the animal awake (12). Mice were single housed in cages with corncob bedding and a handful of soft bedding for nest building for 7 days prior to SD or non-sleep deprivation (NSD). Mice had *ad libitum* access to food and water in these housing conditions and during the sleep deprivation experiments. Mice were habituated to the experimenter and gentle handling stimuli, which was done by the experimenter holding each mouse in the palm for 2min and cage tapping for 2min for 4 consecutive days prior to the 1^st^ sleep deprivation bout. After 5 hours, mice were returned to the original standard housing room where NSD mice were kept undisturbed throughout the experimental time window (ZT0-ZT5). The second SD or NSD bout was performed a week later using the same gentle handling method and timeline. After the second bout of SD/NSD, mice went through perfusion fixation for immunohistochemistry validation of c-Fos/RiboTag expression or cervical dislocation to collect tissue for RNA extraction from the hippocampal CA1.

For active neuron tagging experiments: injection habituation occurred for 2 consecutive days prior to the first sleep deprivation experiment in which mice received an i.p. injection of saline after the 2min of handling and before being placed back in the cage for 2min of light cage tapping. All mice received an i.p. injection of 15mg/kg or 50mg/kg 4-hydroxytamoxifen (4-OHT) at ZT2 on the day of first SD/NSD.

### Adeno-associated virus (AAV) constructs and stereotactic surgeries

Surgeries were performed under isoflurane anesthesia (5% induction, 2% maintenance) following administration of Meloxicam (5mg/kg subcutaneous, s.c.). Animal health was monitored for 5 days following surgery along with administration of Meloxicam (5mg/kg, s.c.) for the first 2 days post-surgery. To label the SD activated hippocampal neurons, 10-12 week old mice were injected intrahippocampally with a cocktail of AAV8-c-Fos-ERT2-Cre-ERT2-PEST-WPRE (titer-1.57E+13 GC/mL, generated and packaged by Stanford University Gene Vector, catalog# GVVC-AAV-139) and AAV9-EF1a.DIO.HA-mRpl22.IRES2.eYFP.WPRE.hGH (titer-1.14E+13 GC/mL, generated and packaged by the University of Pennsylvania Viral Vector Core). For labeling cells activated by SD throughout the hippocampal subregions, the AAV cocktail was injected into the dorsal hippocampus bilaterally (1000nL at a rate of 200nL/min) using the following coordinates relative to bregma: anteroposterior (AP) −1.9mm, mediolateral (ML) ±1.5mm, dorsoventral (DV) −1.5mm from the surface of the brain. For labeling cells activated by SD specifically in area CA1, the AAV cocktail was injected into dorsal CA1 bilaterally (300nL at a rate of 50nL/min), the following coordinates were used upon validation: AP −1.8mm, ML ±1.45mm, DV −1.65 from the surface of the skull. Experiments were performed after 3 to 4 weeks to allow recovery from surgery and for spread of the viral particles in the targeted brain regions.

### Drug preparation

4-OHT solution was prepared fresh on the day of experiment as previously described (34). 4-OHT (Catalog #H6278, Sigma-Aldrich, MO, USA) was dissolved in 100% ethanol at 37°C for 15min with constant rotation to prepare a 20mg/mL stock solution. Stock solution was then diluted to 10mg/mL in 100% corn oil followed by another 15min incubation at 37°C with constant rotation. The solution was vacuum centrifuged for 15min to evaporate the ethanol, yielding an injectable solution of 10mg/mL 4-OHT in corn oil, and was administered intraperitoneally (i.p.) at a dose of 15mg/kg or 50mg/kg body weight.

### Immunohistochemistry (IHC)

Perfusion fixed brains (4% v/v paraformaldehyde in 1X phosphate buffer saline (PBS), Catalog #15710, Electron Microscopy Sciences, PA, USA) were equilibrated in sucrose solution (30%w/v in 1X PBS) for 48h. The brains were then sliced into 30µm sections using a Leica cryostat (Leica CM3050S, IL, USA) at −20°C and stored in cryoprotectant solution (30% w/v sucrose, 30% v/v ethylene glycol, and 0.01% Sodium Azide in 1X PBS). Sections containing the dorsal hippocampus were rinsed three times for 5min each in 1X PBS, followed by a 1h incubation in blocking solution (3% normal Donkey/Goat serum in 0.4% triton-X100/PBS buffer) and an overnight incubation at room temperature (RT) with primary antibodies: rabbit anti-c-Fos (1:3000, #226008, Synaptic Systems, Goettingen, Germany), guinea pig anti-c-Fos (1:3000, #226308, Synaptic Systems), rabbit anti-HAtag (1:2000, #3724, Cell Signaling, MA, USA), mouse anti-NeuN (1:1500, #ab104224, Abcam, MA, USA), rabbit anti-Sox9 (1:1500, #ab185966, Abcam), mouse anti-GAD67 (1:1000, #MAB5406, Sigma-Aldrich). Following overnight incubation with primary antibodies, slices were rinsed three times for 5 min in 1X PBS and incubated with secondary antibodies: Alexa Fluor 488 goat anti-rabbit (1:1000, #A11008, Invitrogen, CA, USA), Alexa Fluor 647 donkey anti-rabbit (1:1000, #A32795, Invitrogen), Alexa Fluor 555 goat anti-guinea pig (1:1000, #150186, Abcam), Alexa Fluor 647 goat anti-guinea pig (1:1000, #A21450, Invitrogen), and Alexa Fluor 488 donkey anti-mouse (1:1000, #A21202, Invitrogen) for 2h at RT followed by three rinses in 1X PBS for 5min each. Slices were mounted onto Superfrost Plus (Fisherbrand) slides with Prolonged Diamond Antifade mountant with DAPI (P36962, Thermo Fisher Scientific, MA, USA) to stain cell nuclei.

### Confocal Microscopy and Image Analysis

Following IHC, images of the dorsal hippocampus were acquired using a Leica SPE Confocal microscope equipped with lasers at 405nm, 488nm, 561nm, and 635nm. All images (8-bit) were obtained with identical settings for laser power, detector gain, and pinhole diameter (1.0AU) using a 20X oil immersion objective at 1024 × 1024-pixel resolution and 1.5X optical zoom.

The Adult Mouse Allen Brain Reference Atlas was used as a reference for the hippocampus and hippocampal subregions. A total of 2-3 sections (2 hippocampi per section) were imaged per mouse and averaged get the number of c-Fos puncta per animal. Images were processed using the ImageJ software. Background fluorescent signals generated from AAV infusion surgery was subtracted from c-Fos and HAtag channels using the Image Calculator function in ImageJ. Images were converted to 16-bit for c-Fos+ cell quantification. The c-Fos+ cells within the cell body layers of each subregion were then plotted and counted automatically using the Analyze Particle function in ImageJ. The HAtag+ cells, DAPI+ nuclei, and double-labeled with HAtag/NeuN/Sox9 cells were counted manually. For c-Fos and GAD67 overlapping comparison, GAD67 images were first merged with the DAPI channel and overlapping cells within cell layers were identified as inhibitory interneurons. Then, interneurons overlapping with c-Fos labeling were quantified as activated inhibitory interneurons in each subregion. The percentage of activated excitatory neuron in each subregion was calculated by subtracting the inhibitory (GAD67+) neuronal fraction from the neuronal (NeuN+) fraction and normalizing to the fraction of NeuN+ neurons.

### Immunoprecipitation and mRNA Extraction

For translatome sequencing, mice were sacrificed at ZT5 by cervical dislocation after the 2^nd^ SD/NSD. Hippocampi were rapidly dissected into ice-cold 1X PBS followed by microdissection of CA1 on ice under the Leica S4E stereo microscope as previously described (35). CA1 tissues were collected in sterile, DNase and RNase-free tubes chilled on dry ice and immediately stored at −80°C. Ribosome-associated mRNA was immunoprecipitated as previously described (29). Both CA1 tissues were binned from 5 mice in each group were homogenized in 1.2mL homogenization buffer (50mM Tris pH 7.4, 100mM KCL, 12mM MgCl_2_, 10% Nonidet P-40 (NP40), 1mM DTT, 200U/mL Promega RNasin, 1mg/mL heparin, 100μg/mL cycloheximide, 1X protease inhibitor (#P8340, Sigma, MA, USA). The homogenates were centrifuged at 10,000x*g* for 10min at 4°C. The supernatant was incubated with 5μL Anti-HA.11 Epitope Tag Antibody (#16B12, Biolegend, CA, USA) for 4h at 4 °C with rotation, followed by overnight incubation with Pierce Protein A/G magnetic beads (#88803, Pierce, MA, USA) at 4 °C with rotation. Supernatant was removed from the beads using a magnetic rack and the beads were washed 3 times with a high salt buffer (50mM Tris pH 7.4, 300mM KCL, 12mM MgCl_2_, 10% NP40, 0.5mM DTT, 100μg/mL cycloheximide). mRNA bound to the beads were eluted at 4°C in 350μL of RLT buffer from the RNeasy Micro Kit (#74004, Qiagen, Venlo, Netherland) supplemented with β-mercaptoethanol mRNA extraction at 10μL/mL. DNAse digestion of IP-eluted mRNA were performed following Qiagen’s RNeasy extraction kit instructions. RNA concentration and quality was assessed using the Qubit^TM^ RNA High Sensitivity Assay Kit (#Q32852, Thermo Fisher Scientific, MA, USA).

### Library Preparation

RNA quality was assessed using an Agilent Bioanalyzer. RNA library preparation from samples of SD and NSD mice were prepared at the Iowa Institute of Human Genetics (IIHG), Genomics Division, using Illumina Stranded Total RNA Prep Ligation with Ribo-Zero Plus v01 sample preparation kit (Illumina, CA, USA). Pooled library was sequenced on the Element Aviti24 sequencer with 150-bp paired-end chemistry at the IIHG core.

### fosTRAP-seq Data Analysis

Data analysis was performed using Partek Flow interface (Partek, Illumina, CA). Following QA/QC, raw reads were subjected to the base trimming to remove poorly scored 3’-bases and aligned using STAR aligner version 2.7.8a (36). Aligned reads were quantified by Partek E/M model and annotated to mouse reference genome mm39 Ensemble release 110. The resulting mRNA counts were filtered to exclude mRNA with minimum count ≤ 5 (1%) and normalized using median ratio method. Differential gene expression analysis was performed with DESeq2 (37). Differentially Expressed Genes (DEGs) were identified by the standard: fold change >1.2 or <-1.2, false discovery rate (FDR) value <0.05.

Enrichment of upregulated/downregulated DEG-associated pathways from the fosTRAP-seq data was performed with a combination of Gene Ontology (GO-Biological Process-EBI-UniProt-GOA-ACAP-ARAP) and Kyoto Encyclopedia of Genes and Genomes (KEGG) database. The GO-BP analyses were performed with the clusterProfiler package in R (38) and the Cytoscape (version 3.10.4 https://cytoscape.org/) (39) plug-in ClueGo (version 2.5.10) (40). Only the common GO-BP pathways between the two methods with an adjusted p value <0.05 and gene counts ≥ 3 were considered as significant. Volcano plot of all DEGs and Cnet plots representing enriched GO-BP pathways with their associated DEGs were generated using SRplot website (41) (Fig. 6). KEGG analyses were performed using the Cytoscape plug-in ClueGo, pathways were considered significantly enriched with an adjusted p value < 0.05 and gene counts ≥ 3. To connect terms in the network, ClueGO utilizes kappa statistics in which here was set as ≥ 0.4 and generated the map figure of enriched functions (additional file1: Fig. S3, 4). For comparison between our fosTRAP-seq and previous excitatory TRAPseq data published by Lisa Lyons et al., (31) (Fig. 6), we first ran the same DEG identification protocol as described above on excitatory TRAPseq raw data and compared common DEGs between the two datasets. The quadrant diagram illustrating common DEGs (Fig. 6) were generated using GraphPad Prism v10. GO-BP analysis was performed to identify enrichment biological processes associated with common DEGs using clusterProfiler package in R (38) and the pathways with their associated genes were presented using a Sankey diagram generated using SRplot (41) as well. Biological functions associates with DEGs specific for each database were performed using KEGG analysis (additional file1: Fig. S4) with Cytoscape as described.

### Statistical analysis

All statistical analyses were performed using GraphPad Prism v10 with α = 0.05 for analyses. Imaging data comparing c-Fos and RiboTag expression and neuronal reactivation were analyzed using unpaired two-tailed t-tests to compare the NSD and SD groups (Fig. 1, 3C, 4, 5) and the distal and proximal CA1 sections (additional file1: Fig. S2). Two-way analysis of variance (ANOVA) comparisons were used to determine the interaction between c-Fos activation and number of sleep deprivation epochs (Fig. 3D, E). Data is presented as Mean ± SEM and *n* represents the number of animals in all experiments, IHC data is presented as an average of 3-4 sections per animal per condition. In figures, ns refers to non-significant, * refers to a *p* value < 0.05, ** refers to *p* < 0.01, *** refers to *p*<0.001, and **** refers to *p*<0.0001.

**Figure 1.**
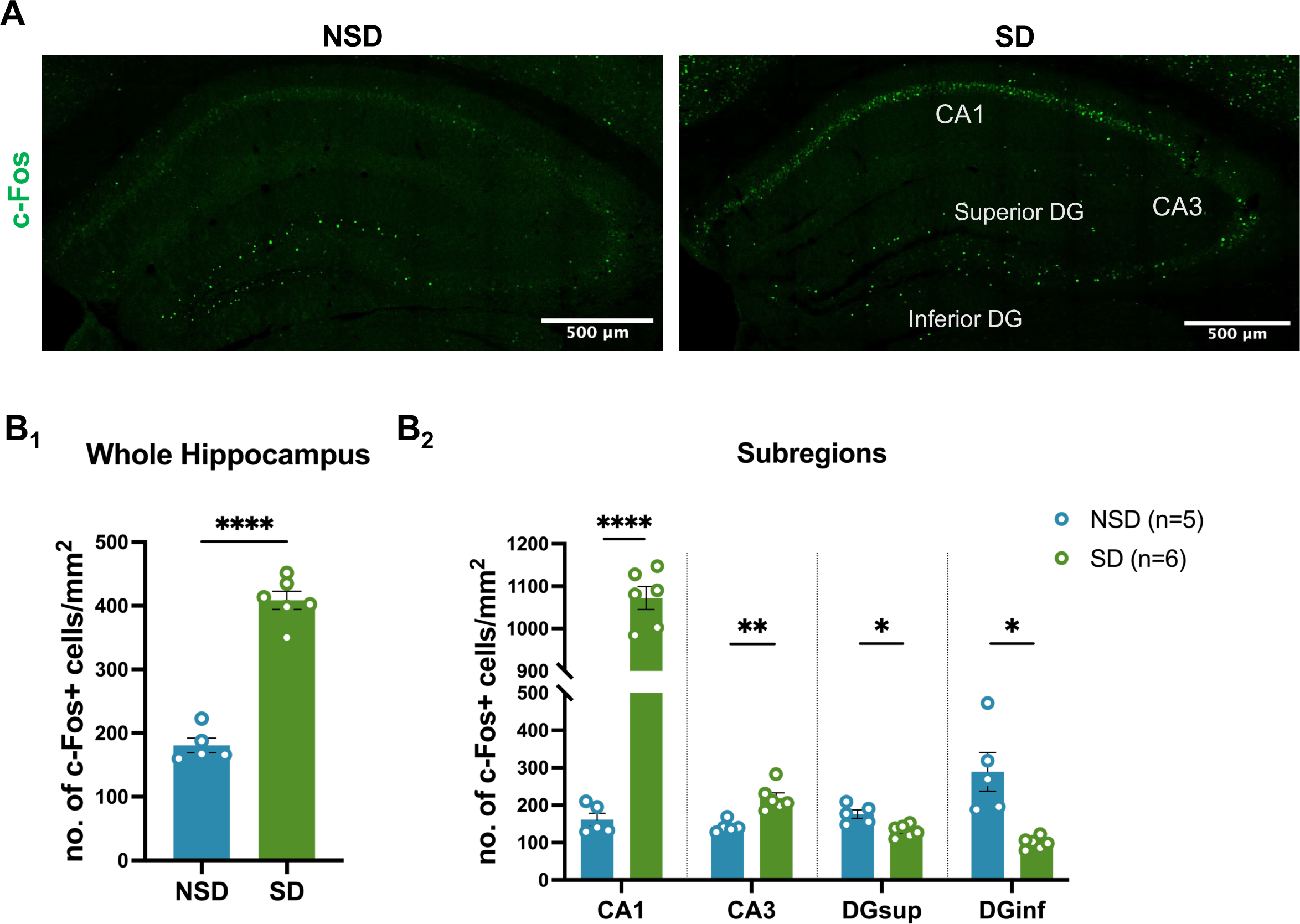
cFos expression is differently altered within hippocampal subregions after acute sleep deprivation. **A**) Immunofluorescent images of c-Fos expression in NSD and SD hippocampus. **B**) Comparison of area normalized cFos expression between NSD (n=5) and SD (n=6) groups from the entire hippocampus (1) and within different subregions (2): CA1, CA3, and superior and inferior blades of Dentate Gyrus (DGsup and DGinf). Data are shown as mean ± standard error of the mean (SEM). Unpaired two-tailed t test for B_1_: NSD=180.64±11.53, SD=408.39±14.27, t(9)=12.05, p<0.0001. Multiple unpaired t tests for B_2_: CA1: NSD=161.50±17.15, SD=1072.02±26.98, t(9)=27.10, p<0.00001; CA3: NSD=142.53±6.97, SD=218.99±14.11, t(9)=4.54, p=0.0014; DGsup: NSD=176.18±11.26, SD=131.76±6.58, t(9)=3.56, p=0.0061; DGinf: NSD=289.11±51.80, SD=99.07±6.38, t(9)=4.02, p=0.0030.

## RESULTS

### Sleep deprivation causes non-uniform activation of hippocampal subregions as measured by c-Fos expression

Induction of IEGs, such as *Fos, Arc, and Egr1*, mark recent neuronal activation, and previous studies have investigated the impact of sleep and sleep loss on neuronal activity across rodent brains by measuring mRNA or protein levels of IEGs (24–27, 30, 32, 42). To investigate the impact of acute SD on neuronal activation within hippocampal subregions, we sleep deprived mice for 5h by the gentle handling method. This method was chosen to avoid confounds with methods that may cause novelty-induced hippocampal activation. Following our 5h SD, we mapped c-Fos protein expression, a well characterized marker of neuronal activity, within the hippocampus as an indicator of neuronal activation. Quantification of c-Fos protein abundance by immunohistochemistry in SD and NSD mice found increased c-Fos in SD samples suggesting an overall increase in hippocampal activity (Fig. 1A and B1) after 5h of SD. As the hippocampal subregions have distinct functions, we quantified c-Fos expression within each hippocampal subregion finding that sleep deprivation for 5h induced c-Fos expression in pyramidal cell layers in CA1 and CA3 subfields. However, we observed decreased c-Fos expression in the superior (DGsup) and inferior blades (DGinf) of the DG (Fig. 1B_2_). Furthermore, although CA1 and CA3 both showed an increase in c-Fos expression after 5h of SD, the induction of c-Fos expression was more prominent in CA1 (8-fold induction compared to NSD) compared to CA3 (1.6-fold induction compared to NSD).

We then asked whether a specific cell type within each subregion is selectively activated by sleep deprivation. To answer that question, we performed co-staining with c-Fos and cell-type specific markers (NeuN for neurons and Sox9 for astrocytes). Our results suggest that neurons (NeuN+) represent the majority of activated cells in all the subregions while a minor fraction of those c-Fos+ cells in each subregion are astrocytes (Sox9+) (Fig. 2A-B). In addition, depending on the subregion, a variable fraction of activated cells (c-Fos+) was classified as other cell types (e.g. microglia, oligodendrocytes) because they were negative for NeuN and Sox9. To further identify whether those c-Fos+ neurons are excitatory or inhibitory, we performed double labeling against c-Fos and GAD67, results indicated that sleep deprivation significantly activates excitatory pyramidal neurons (about 95% of all activated neurons) rather than inhibitory interneurons in all hippocampal subregions (Fig. 2C-D).

**Figure 2.**
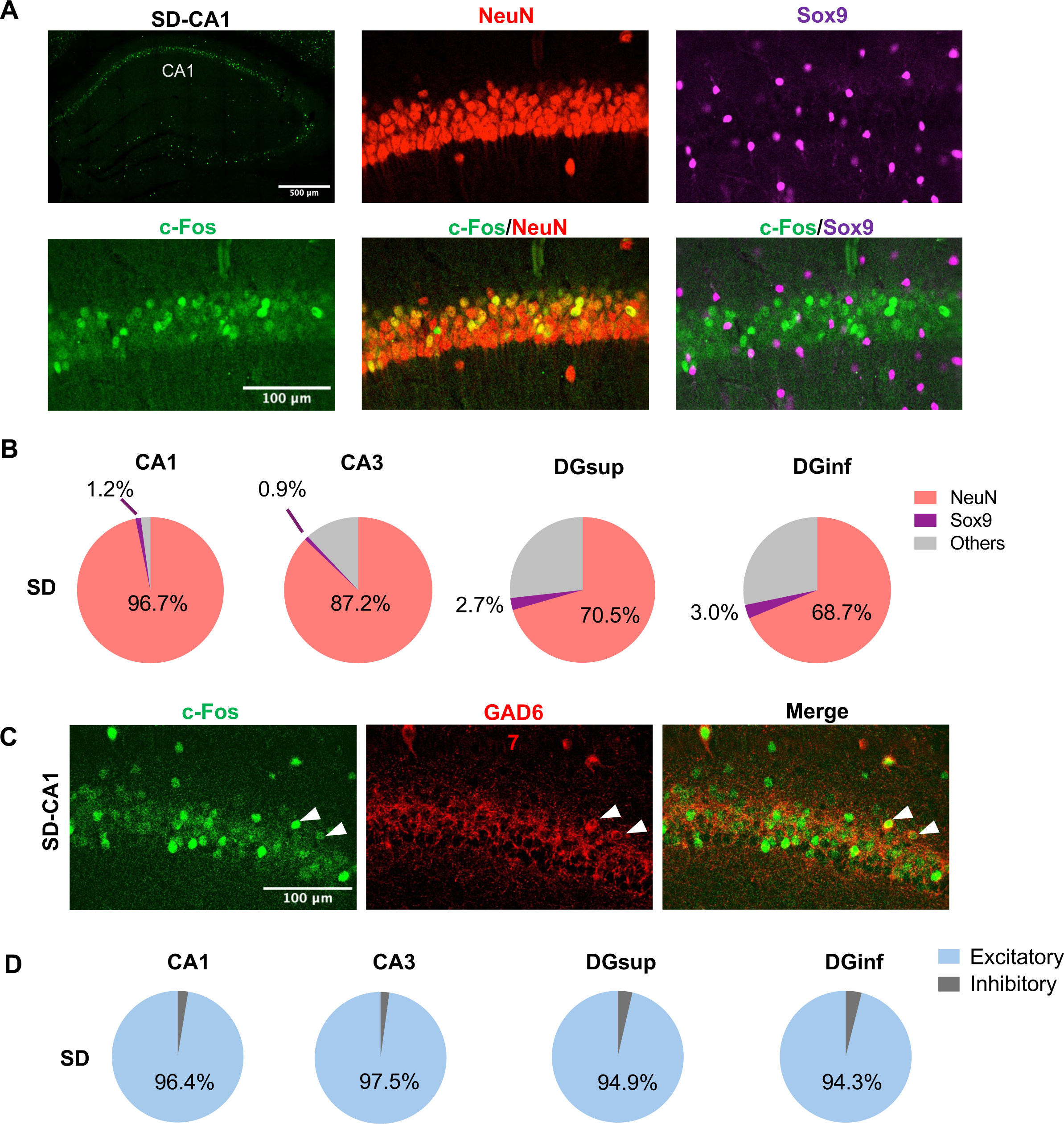
Excitatory neurons are selectively activated by SD. **A**) Representative images of c-Fos double labeling with NeuN and Sox9 after SD. **B**) Percentage of c-Fos+NeuN+ and c-Fos+Sox9+ double labeled cells to the total number of c-Fos+ cells within each subregion in SD (n=3) hippocampus. **C**) Representative images of c-Fos double labeling with GAD67 (inhibitory neuronal marker) after SD. **D**) Percentage of excitatory and inhibitory neurons to the neuronal population within cell layers of each subregion in SD (n=3) hippocampus.

### Repeated SD has similar effects on c-Fos expression

In humans, SD is not usually a singular event, but frequently occurs on multiple nights or a weekly basis. However, how hippocampal neuronal activity is affected by repeated sleep loss remains poorly understood. To address this gap and to determine whether the prominent c-Fos induction in CA1 was due to stochastic neuronal activity or an inherent bias that makes a subset of neurons susceptible to sleep loss, we subjected animals to a repeated SD paradigm (Fig. 3A). In this paradigm animals were subject to SD twice with a 1 week-long recovery between the first and the second SD. This allowed us to determine how repeated SD alters c-Fos expression across hippocampal subregions. A second SD period revealed a similar pattern of c-Fos expression in the hippocampus as a single SD with induction in CA1 and CA3, and a decreasing trend in DG inferior blade (Fig. 3 B, C). The repeated protocol itself does not have a significant effect on c-Fos abundance within subregions. We observed similar c-Fos expression levels between single and repeated NSD and SD groups (1xNSD vs 2xNSD, and 1xSD vs 2xSD) within subregions, although the DG region showed decreased c-Fos after repeated NSD (Fig. 3D, E & additional file2: Table S1). These results indicate consistent levels of neuronal activation after each individual SD session. More importantly, the c-Fos induction patterns evident with a single SD were conserved after the second SD: high c-Fos induction in CA1 (7-fold) and a smaller induction in CA3 (3.5-fold).

**Figure 3.**
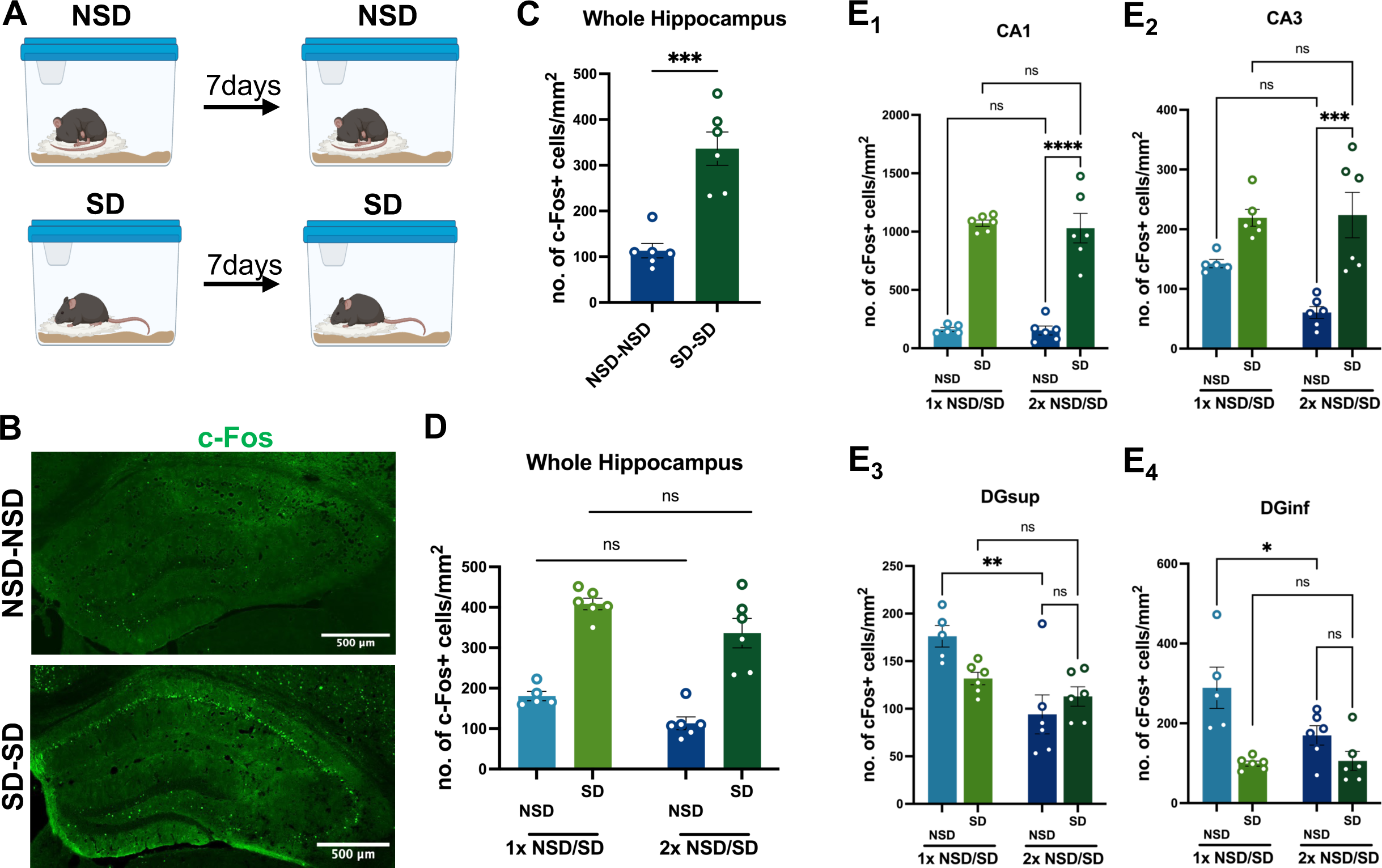
Repeated sleep deprivation induces similar pattern of cFos expression in the hippocampus. **A**) Schematic of repeated sleep deprivation. **B**) Immunofluorescent images of c-Fos expression after repeated NSD (NSD-NSD) and repeated SD (SD-SD) in the hippocampus. **C**) Comparison of area normalized c-Fos expression between NSD-NSD (n=6) and SD-SD (n=6) whole hippocampus. Unpaired two-tailed t-test: NSD=113.13±15.84, SD=336.38±36.42, t(10)=5.62, p=0.0002. **D, E**) Comparison of area normalized cFos expression between single (1x NSD/SD) and repeated sleep deprived (2x NSD/SD) hippocampus (D) and hippocampal subregions (E). Data are shown as mean ± SEM. D: Two-way ANOVA shows a significant effect of sleep deprivation (F(1,19)=98.43, *p*<0.0001) and a significant effect of repeat manipulation (F(1,19)=9.42, *p*=0.0063), but no significant effect of interaction (F(1,19)=0.01, *p*=0.9224). E: Two-way ANOVA for each subregion: E1: effect of SD (F(1,19)=158.5, p<0.0001), ns for effect of repeat manipulation and interaction; E2: effect of SD (F(1,19)=29.29, p<0.0001), ns for effect of repeat manipulation and effect of interaction; E3: ns for effect of SD, effect of repeat manipulation (F(1,19)=14.23, p=0.0013), effect of interaction (F(1,19)=5.57, p=0.0291); E4: effect of SD (F(1,19)=19.45, p=0.0003), ns for effect of repeat manipulation, effect of interaction (F(1,19)=4.81, p=0.041). Bonferroni post hoc tests were performed for panel D & E to compare single vs repeated exposure under SD & NSD conditions, as well as repeated NSD vs SD c-Fos expression. Detailed 2-way ANOVA analyses results see additional file2: Table S1.

Together, the results suggest that SD consistently induces region-specific changes in c-Fos expression and neuronal activation within the hippocampus, with the CA1 and CA3 regions showing increased c-Fos and the DG consistently showing decreased c-Fos expression.

### c-Fos driven RiboTag labels hippocampal neurons activated by sleep deprivation

Although there is a significant induction of c-Fos expression in CA1 pyramidal neurons after both single and repeated SD compared to NSD, the number of c-Fos+ neurons represent a small portion of the total number of CA1 pyramidal neurons. While the results from the repeated SD paradigm suggest that CA1 neurons are activated by SD, it remains unclear if the neurons within CA1 that show c-Fos induction are randomly recruited by individual SD sessions or whether neurons activated by an initial SD period are then more likely to be activated by additional periods of SD. To further characterize the neurons activated by repeated bouts of SD, we took advantage of the fosTRAP system, which can label neurons activated by defined stimuli (28).

We employed an AAV-based neuron labeling system in which animals were injected intrahippocampally with a cocktail of a c-Fos driven Tamoxifen-dependent Cre vector (AAV8-Fos-ERT2-Cre-ERT2) and a Cre-dependent RiboTag vector (AAV9-EF1a-DIO-HA-mRpl22-IRES-eYFP). Administration of 4-OHT facilitates expression of the tamoxifen-inducible c-Fos-driven Cre recombinase (Cre^ERT2^) upon neuronal activation, which subsequently drives expression of mRpl22-HA (RiboTag) (28, 29) (Fig. 4A) to label CA1 pyramidal neurons. Using this strategy, we first tested CA1 neuronal labeling efficacy, indicated by RiboTag (HAtag) expression, using different labeling durations, 4-OHT doses, and AAV infusion methods (additional file1: Fig. S1 and Fig. 4).

**Figure 4.**
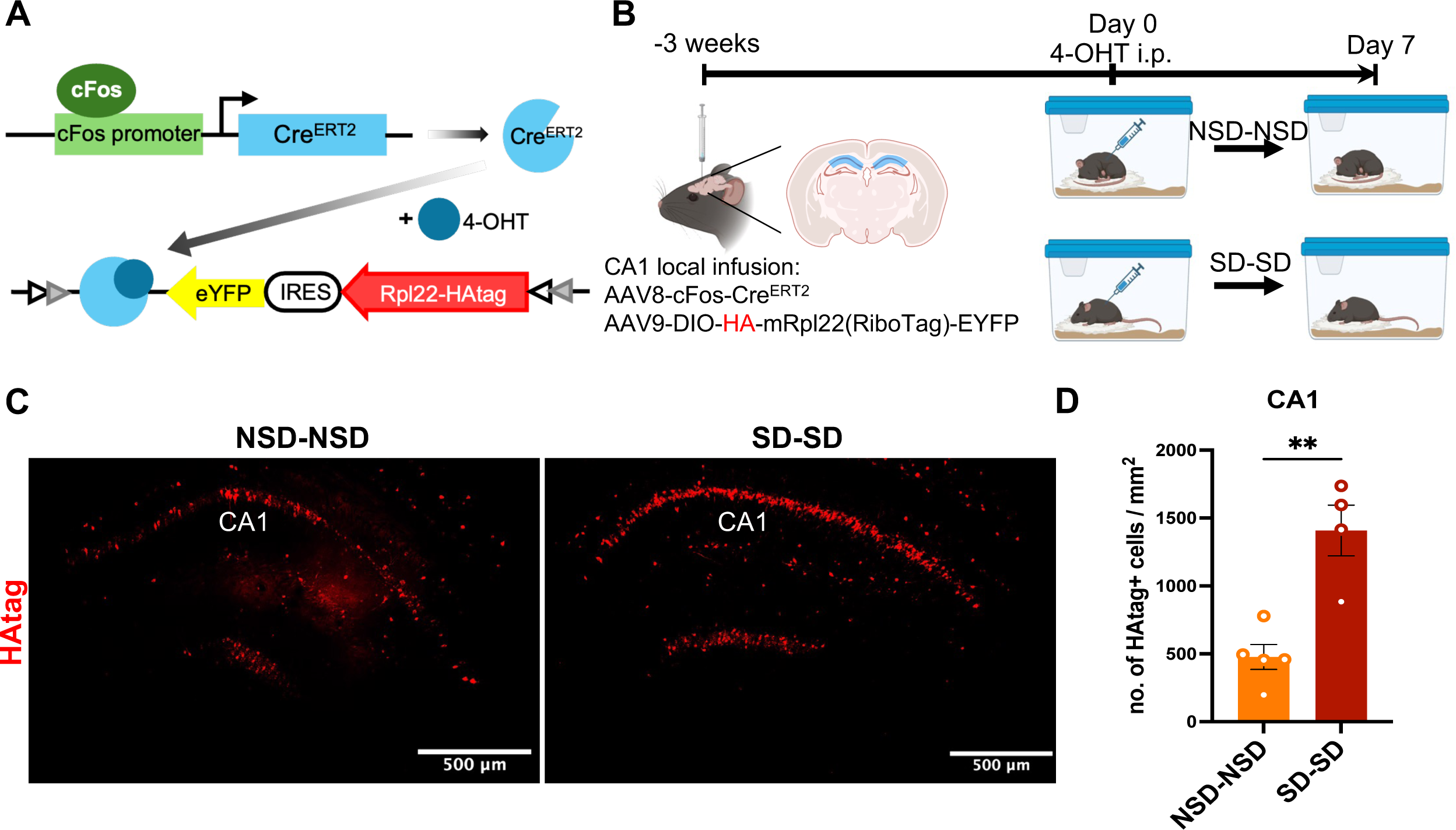
cFos-RiboTag strategy to label activated cells in area CA1. **A)** Diagram of the activity (cFos)-driven ribosomal tagging (RiboTag) strategy**. B**) Diagram of CA1 neuron labeling using cFos-RiboTag strategy with repeat sleep deprivation. **C**) Representative images of neuron labeling shown as RiboTag (HAtag) expression in the NSD and SD hippocampus. **D**) Neuronal labeling efficacy in hippocampal CA1: area normalized HAtag+ cells in NSD (n=5) and SD (n=4) groups. Unpaired two-tailed t-test performed, NSD=476.91±02.01, SD=1408.06±186.79, t(7)=4.79, p=0.0020. Data shown as mean ± SEM.

Unlike c-Fos protein itself, which is expressed rapidly following neuronal activation, expression of the c-Fos activation-promoted RiboTag depends on Cre^ERT2^ mediated recombination of loxP sites and consequent expression of mRpl22-HA (RiboTag). We first assessed the amount of time needed to get robust expression of HA following 4-OHT delivery and SD or NSD. In a 5h labeling test (additional file1: Fig. S1A-C), where 4-OHT was administered at ZT0 and tissue was collected after SD or NSD (ZT0-5), we observed only weak expression of the HAtag, and there were no differences between the SD and NSD groups, confirming that the RiboTag marker takes longer to develop. To provide a longer expression window, we waited one week based on previous results showing that repeated sleep deprivation leads to c-Fos activation in CA1 and CA3 that are comparable to a single SD (Fig. 3). Consequently, this approach requires the use of c-Fos-driven RiboTag labeling for neurons activated by an initial period of SD and subsequent c-Fos immunohistochemistry to identify neurons activated by a second period of sleep deprivation. We divided mice into repeated sleep deprivation (SD-SD) or repeated sleep (NSD-NSD) groups. All mice received an injection of 4-OHT at ZT2 of the 1^st^ SD/NSD (34) to drive RiboTag expression in neurons activated during the 5h of SD/NSD. We also tested two concentrations (15mg/kg and 50mg/kg) of tamoxifen and found that a higher dose of 4-OHT (50mg/kg) was necessary to reflect the proportion of cells activated by SD (additional file1: Fig. S1D-F).

To further optimize RiboTag expression, we targeted the AAV cocktail infusion to hippocampal area CA1 bilaterally. Specific CA1 expression revealed a 3-fold induction of RiboTag expression after SD compared to the NSD group (Fig. 4). The expression level of RiboTag using this CA1 targeting strategy was higher than the normalized number of cells showing c-Fos expression that we observed in both NSD and SD for the single or repeated SD groups (Fig. 1 and 3), particularly for the NSD groups. This could be due to the short half-life (∼2h) of c-Fos (43), and consequently the c-Fos protein that was induced in the early stages of the light cycle may not be reliably detectable at ZT5 using IHC in both the NSD and SD conditions, whereas the RiboTag will persist upon neuronal activation. Another factor that is likely to increase RiboTag expression in the NSD group is the unavoidable wakefulness and subsequent c-Fos expression induced by i.p. injection of 4-OHT. Together, these optimization strategies established a method for activity-driven RiboTagging that successfully labels CA1 pyramidal neurons that are activated by SD.

### CA1 pyramidal neurons are reactivated by repeated SD

To answer the previous question that if subsets of neurons reactivate during repeated SD in the CA1 or whether hippocampal CA1 pyramidal neurons are randomly activated by repeated sleep loss, we performed RiboTagging combined with c-Fos immunolabeling. Immunofluorescence was used to identify the cells positive for RiboTag (the neurons that were activated during first SD or NSD) and c-Fos (the neurons that were activated during the second SD or NSD). Cells that are positive for both c-Fos and the HA tag are neurons that undergo reactivation during the second SD period (Fig. 5A). Our analysis showed that about 30% of CA1 neurons activated by the first SD (HAtag positive) were reactivated by the second SD (c-Fos positive; Fig. 5B). Furthermore, of the CA1 pyramidal neurons activated by the second SD (c-Fos+), 40% were previously activated by the first SD (Fig. 5C). We examined HAtag and c-Fos immunolabeling across CA1 and found higher HAtag and c-Fos expression in proximal CA1, where pyramidal neurons are mainly driven by medial entorhinal cortex projections and involved in spatial information processing (44, 45), with similar reactivation rate between proximal and distal CA1 (additional file1: Fig. S2), which aligns with impaired spatial memory.

**Figure 5.**
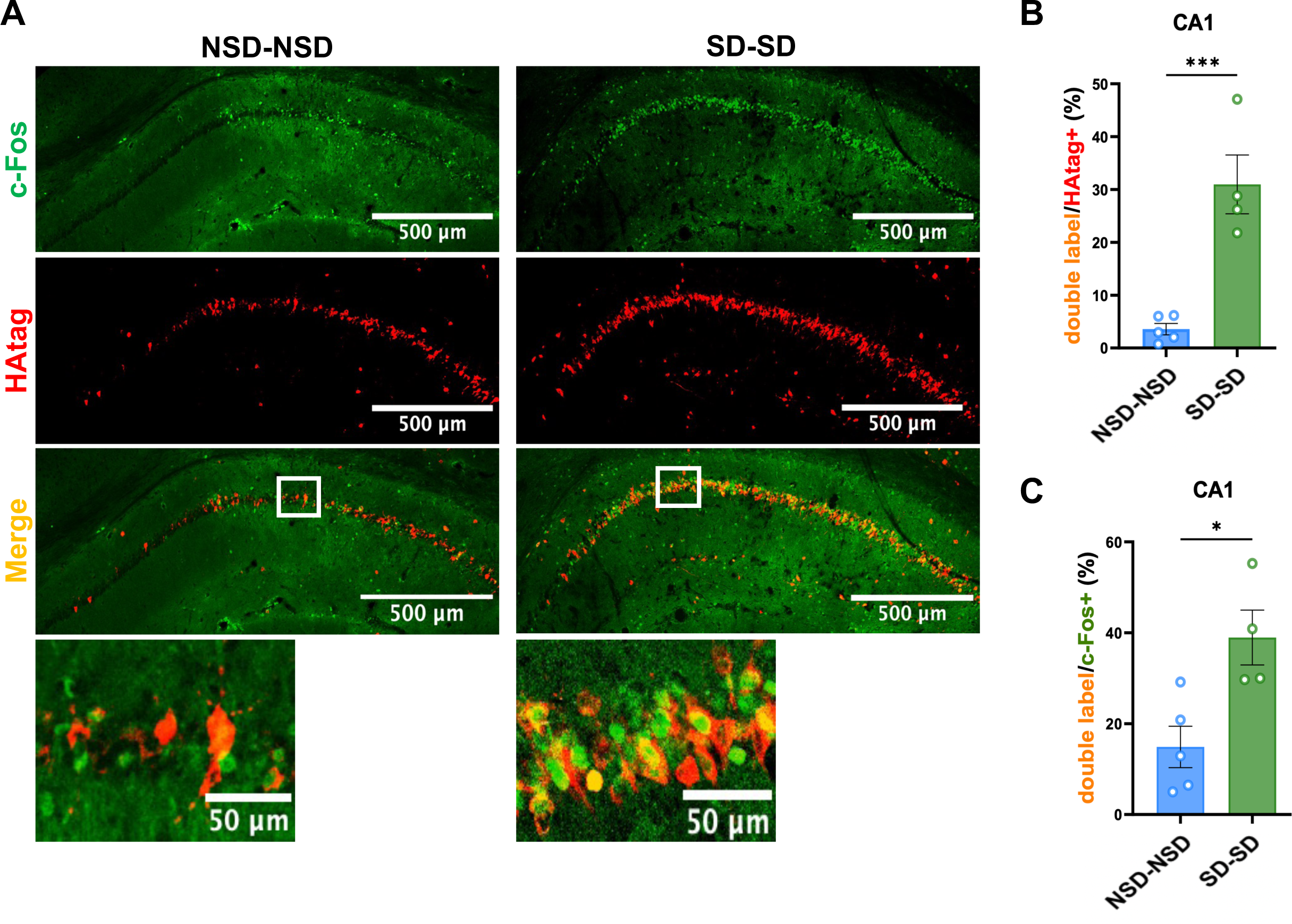
CA1 pyramidal neurons are reactivated by repeated SD. **A**) Immunofluorescence images of cFos and HAtag double labeling in area CA1 in NSD-NSD and SD-SD groups. **B, C**) Neuronal reactivation: percentage (%) of double labeled cells / HAtag+ cells (B) and double labeled cells / cFos+ cells (C) overlap in NSD-NSD (n=5) and SD-SD (n=4) CA1. Unpaired two-tailed t-test. Data shown as mean ± SEM. B: NSD-NSD=8.82±2.71, SD-SD=30.97±5.56, t(7)=5.44, p=0.0064. C: NSD-NSD=36.02±7.95, SD-SD=38.96±6.03, t(7)=3.26, p=0.7866.

Importantly, overlap in c-Fos and HAtag was minimal in the NSD-NSD group: about 5% of RiboTagged cells during 1^st^ NSD were activated by the 2^nd^ NSD (Fig. 5B), and double-labeled cells make up approximately 15% of the limited population of c-Fos+ cells after the second NSD (Fig. 5C). This overlap is likely due to background RiboTag expression (additional file1: Fig. S1G). This neuronal reactivation confirms that repeated SD stimulates a similar population of pyramidal neurons in hippocampal CA1 rather than a random activation, which suggests that a subset of neurons is more likely to be activated during single and repeated SD.

### Activity-dependent translatome changes in SD-sensitive CA1 neurons

In addition to tagging activated neurons, the RiboTag approach we applied also allows analyzing gene expression and mRNA translation in specific cell types (29, 31), in our case the c-Fos+ neurons, by performing Translating Ribosome Affinity Purification with RNA sequencing (fosTRAP-seq). We tagged SD-sensitive CA1 neurons by subjecting all mice to sleep deprivation on day 0, followed by division into NSD and SD groups on day 7 to induce changes in activity and gene expression (Fig. 6A). Immediately after that SD/NSD session, actively translating mRNA from CA1 RiboTag-labeled neurons were pooled using immunoprecipitation with antibodies for the HAtag. Following TRAP RNA extraction (5 animals pooled per sample), we used an unbiased RNA sequencing approach, aligned raw reads using STAR aligner (36), and identified differentially expressed genes (DEGs) by the criterion of: FDR < 0.05, and fold change >1.2 or <-1.2 to avoid false positives. From our fosTRAP-seq data, we identified 631 DEGs being upregulated while 221 DEGs being downregulated in RiboTag labeled CA1 neurons (Fig. 6B, additional file2: Table S2). As expected, the IEGs (such as *Fosb, Arc, Egr1, and Homer1*) that are bound for translation are upregulated by sleep deprivation indicating reactivation of certain SD-sensitive CA1 neurons during the repeated sleep loss. We also observed decreased ribosome associated transcripts of *Cirbp, Rbm3*, *Srsf5* which have been reported by previous bulk transcription analysis (33, 46).

**Figure 6.**
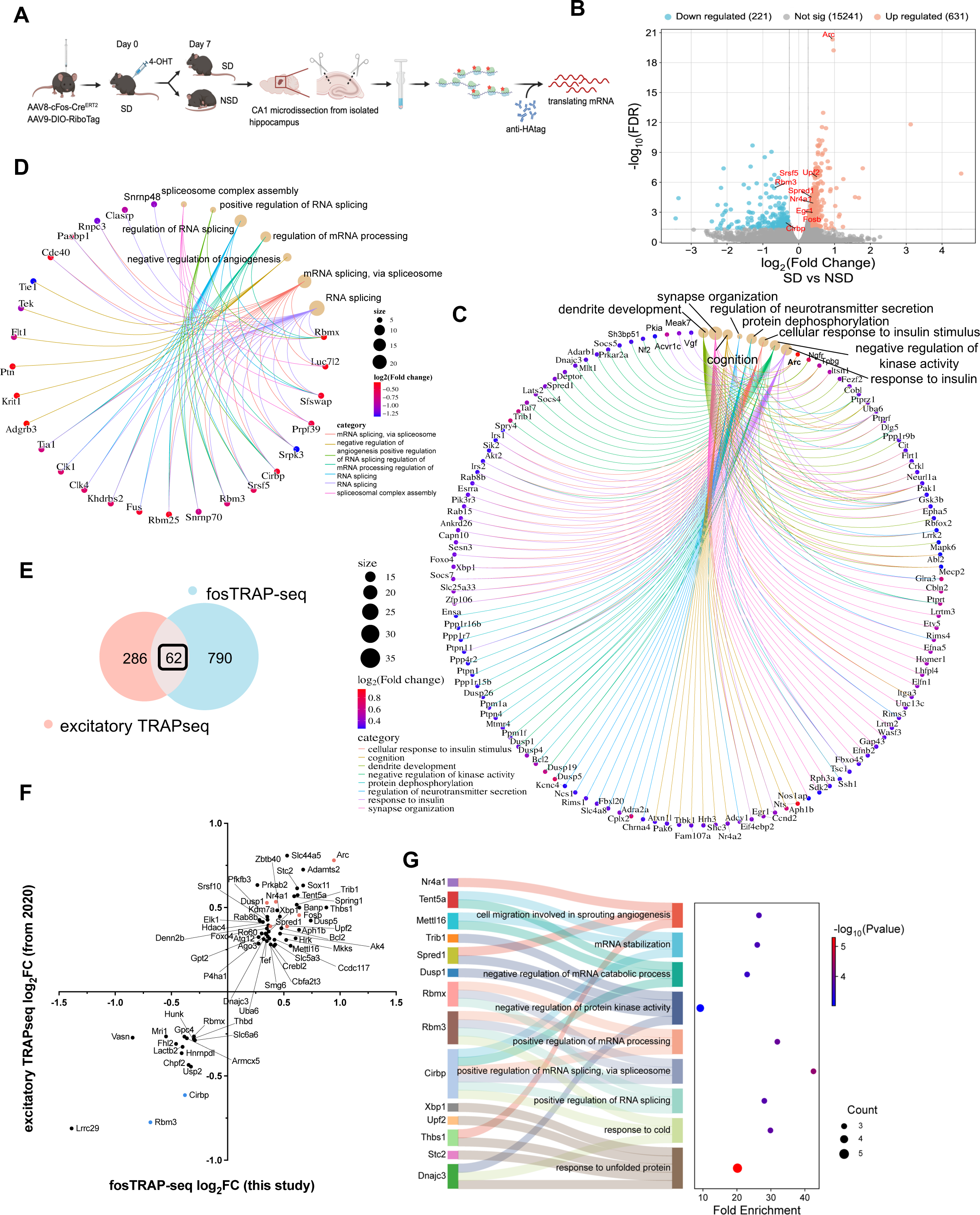
fosTRAP-seq reveals activity-dependent gene expression alteration in SD-sensitive hippocampal CA1 neurons after sleep deprivation. **A)** Diagram of translating mRNA extraction from RiboTag labeled CA1 neurons. All mice were sleep deprived on day 0 then randomly divided into NSD and SD groups on day 7. (n=10 mice/group, CA1 from 5 mice were pooled together for further RNA extraction.) **B**) Volcano plot of all DEGs (FDR<0.05, |FC|>1.2) identified in response to sleep deprivation in hippocampal CA1 SD-sensitive neurons. **C, D**) Cnet plot showing GO biological process (BP) analyzed pathway enrichment (adjusted Pvalue<0.05, gene≥3) associated with DEGs significantly upregulated (C) or downregulated (D) after sleep deprivation in SD-sensitive CA1 neurons. Full details of C, D see additional file2: Table S3. **E**) Venn diagram representing the number of DEGs overlapped between our fosTRAP-seq results and our previously published (Lyons et al., 2020) excitatory TRAPseq after SD identified 62 common genes, 790 only in fosTRAP-seq and 286 only in excitatory TRAPseq. **F**) Quadrant diagram showing comparison of common DEGs between fosTRAP-seq and excitatory TRAPseq. **G**) Sankey diagram (left) shows association of overlapped DEGs to the identified GO biological processes (p<0.05), along with pathway enrichment values as a Dot plot (right). Full details of G see additional file2: Table S6.

We then separately analyzed the upregulated and downregulated transcripts for the enriched biological processes and functions that were affected by SD against the GO: biological process (GO-BP) and KEGG databases. The analyses on the upregulated DEGs revealed strong enrichment in biological processes of protein dephosphorylation (e.g. *Dusp1/4/5/19/26, Ppm1a/1f, Ptpn1/4/11*), negative regulation of kinase activity (e.g. *Adarb1, Dnajc3, Pkia, Dusp1/19*), response to insulin (e.g. *Akt2, Foxo4, Gsk3b, Irs1/2, Capn10, Egr1, Tsc1, Vgf, Ptpn1/11*), dendrite development (e.g. *Arc, Cobl, Mack6, Mecp2, Ngfr, Ppp1r9b*), regulation of neurotransmitter secretion (e.g. *Adra2, Cplx2, Kcnc4, Lrrk2, Rim3/4*), cognition (e.g. *Arc, Egr1, Ccnd2, Nr4a2, Uba6*) having higher enrichment scores among all altered pathways (Fig. 6C, additional file2: Table S3-1), and the biological functions of longevity regulation and insulin signaling pathways (additional file1, 2: Fig. S3A, Table S4-1). The downregulated DEGs are mainly associated with regulation of mRNA processing and splicing processes (e.g. *Cirbp, Rbm3/x, Srsf5, Luc7l2, Cdc40, and Sfswap*), as well as negative regulation of angiogenesis (*Adgrb3, Flt1, Krit1, Ptn, Tek, and Tie1*) (Fig. 6D, additional file2: Table S3-2), and the biological functions of ECM-receptor interaction and spliceosome (additional file1,2: Fig. S3B, Table S4-2).

We also compared these data to our previous work performing TRAPseq (31) on bulk hippocampal excitatory neurons from the hippocampus after 5h of sleep deprivation. There are 62 DEGs overlapped between our fosTRAP-seq and the excitatory TRAPseq and all DEGs share the same direction of alteration (Fig. 6E-F, additional file2: Table S5). Among the common DEGs, IEGs (including *Fosb, Arc, Homer1, Nr4a1*), *Upf* 2 (regulator of nonsense mediated mRNA decay), and *Spred1* (negative regulation of kinase activity/protein phosphorylation) are upregulated in both datasets. The *Cirbp* and *Rbm3* genes, involved in RNA splicing and mRNA processing regulation, are downregulated from the two datasets. GO-BP analysis indicates that protein phosphorylation, mRNA splicing and processing processes are shared in both datasets (Fig. 6G, additional file2: Table S6). We also compared distinct DEGs between the two datasets and KEGG analysis revealed that biological functions of insulin-signaling pathway, longevity regulation pathway, and focal adhesion being enriched specifically in our fosTRAP-seq, while melanogenesis and apoptosis are altered only in excitatory TRAPseq (additional file1,2: Fig. S4, Table S7). The shared and unique DEGs between the two datasets further validate the reliability and robustness of our fosTRAP-seq method.

Taken together, our fosTRAP-seq identified activity-dependent translatome alteration in SD-vulnerable CA1 neurons with a higher specificity and resolution revealing post-transcriptomic-and translational-modification, synapse connection, neuron development, insulin signaling being vulnerable functions affected by sleep loss.

## DISCUSSION

The hippocampus is a structurally distinct and functionally diverse brain region, comprised of interconnected subregions that contribute uniquely to memory encoding processes (7–9). To better understand how sleep deprivation affects hippocampal function, we investigated c-Fos expression patterns across different hippocampal subregions. In this study, we sleep-deprived animals from ZT0-ZT5 by gentle handling, a widely accepted approach that minimizes stress (47) while avoiding the confounding effects of spatial or object novelty on neuronal activation in the hippocampus (47, 48), and examined hippocampal subregional c-Fos expression after single and repeated SD. Results indicated that c-Fos expression is altered differently among hippocampal subregions: CA1, particularly proximal CA1, shows the most robust c-Fos induction; CA3 shows a moderate activity increase; and the DG shows reduced c-Fos expression. (Fig. 1-3). Using the c-Fos driven RiboTag labeling system, we observed 30% to 40% overlap ratio between neurons activated during the first and second SD and confirmed that CA1 pyramidal neurons are reactivated by repeated sleep loss (Fig. 4-5). We then performed fosTRAP-seq and revealed translatome changes in RiboTag labeled CA1 neurons that are sensitive to sleep deprivation (Fig. 6).

In the present study, we observed distinct patterns of neuronal activation among hippocampal subregions caused by sleep deprivation. Hippocampal neuronal activation is regulated by coordinated interactions among glutamatergic, GABAergic, serotonergic, and noradrenergic systems, and regional differences in receptor expression may account for distinct c-Fos activation patterns. During sleep deprivation, there are a multitude of changes in neurotransmitter systems and signaling that differ between areas of the cornu ammonis and dentate gyrus (49, 50). Of note, locus coeruleus–derived norepinephrine release is elevated during SD and is known to induce c-Fos expression and enhance CA1 pyramidal neuron excitability via β-adrenergic signaling (51–53). Consistent with this, our fosTRAP-seq data show that CA1 neurons are enriched for *Adrb1* relative to *Adrb2* and *Adrb3*, suggesting elevated β_1_-adrenergic receptor expression enhances sensitivity to norepinephrine and promotes c-Fos activation via cAMP-CREB signaling. This differential pattern of neuronal activation is consistent with previous spatial transcriptomic studies, which identified unique gene expression changes and enriched molecular functions specific to hippocampal subregions after SD (49, 54) and adds to a growing literature emphasizing the heterogeneity of SD effects in various hippocampal subregions. However, our findings differ from a recent study by Sarma and colleagues (2025), which reported that 3h, but not 6h, of gentle handling sleep deprivation alters c-Fos expression across hippocampal subregions (55). In contrast, our 5 h sleep deprivation paradigm produced a more robust increase in c-Fos, particularly in CA1, which may reflect the higher sensitivity of immunofluorescence compared with DAB-based immunohistochemistry. Notably, our c-Fos expression pattern is consistent with prior genomic, translational, and behavioral studies showing upregulation of c-Fos and other IEGs after 5 h of sleep deprivation (31, 33, 46). This same 5 h gentle-handling protocol reliably induces hippocampus-dependent memory deficits and synaptic plasticity alterations (12, 14) without elevating corticosterone or confounding behavioral stress effects (47, 48). Together, these findings suggest that sleep-loss duration is a critical determinant of hippocampal c-Fos responses, with 5 h representing a key time point at which molecular, electrophysiological, and behavioral consequences of sleep deprivation converge.

Utilizing a viral c-Fos-RiboTag active neuron tagging approach (28, 29), we successfully captured the robust induction of CA1 neuronal activation and identified a specific population of CA1 pyramidal neurons that is repeatedly activated by sleep deprivation. Our approach labeled a substantial number of activated neurons compared to a previous study that reported minimal labeling of CA1 neurons following sleep deprivation using a TRAP transgenic mouse model (56). Our results also demonstrate over 30% overlap between c-Fos+ and RiboTag-labeled neurons across repeated sleep deprivation events (Fig. 5). This represents a significant overlap of neuronal activation after repeated SD comparable to the hippocampal engram reactivation rate observed using IEG-dependent tagging methods (e.g. TRAP2 and Tet-tag) during memory consolidation and retrieval, which typically ranges from 5-20% (49, 57–59). This prominent reactivation in our dataset provides compelling evidence supporting the hypothesis that a specific subset of CA1 pyramidal neurons have distinct molecular or electrophysiological characteristics (60, 61) that confer heightened sensitivity to repeated and chronic sleep loss. Notably, c-Fos expression is widely used as indicator of engram activation and reactivation in hippocampus-dependent memory formation and consolidation. In CA1, higher neuronal place field preference, place field activity, stability of spatial maps, and more precise spatial information decoding is found in the neurons that were c-Fos positive during spatial learning tasks (62). Thus, their strong activation during sleep deprivation –despite the absence of learning– raises a mechanistic question regarding the functional significance of CA1 activation under sleep loss. Consistent with this idea, several hours of sleep deprivation following learning similarly induces CA1 c-Fos expression yet impairs memory consolidation (49, 55), implying that sleep loss may either recruit neuronal populations distinct from memory engrams or aberrantly activate learning-related ensembles while disrupting gene expression, protein synthesis, or sharp-wave ripple–mediated replay (63–65). Future experiments comparing whether learning and sleep deprivation activates a similar population of CA1 pyramidal neurons could be helpful answer the question. Together, these observations highlight a sleep deprivation-sensitive CA1 neuronal subpopulation and provide a framework for dissecting how aberrant neuronal activation and reactivation contribute to hippocampal dysfunction and memory impairment under chronic sleep loss.

Taking advantage of the c-Fos-RiboTag system combined with repeated SD paradigm, we isolated activity-dependent translatome changes in CA1 neurons that are selectively sensitive to sleep loss (c-Fos+). Our fosTRAP-seq analysis identified DEGs that have been reported in previous gene expression studies after acute sleep deprivation (31, 33, 46, 54) supporting the robustness of our repeated SD and neuronal labeling strategy. We reported upregulated translation of transcripts involved in synaptic vesicle exocytosis (e.g. *Cplx2, Rims4, Slc4a8*), regulation of synaptic plasticity (e.g., *Arc, Egr1, Adcy1/8*), and dendrite development (e.g. *Arc, Cobl, Mack6, Mecp2*) all of which are essential for memory consolidation. In contrast, transcripts associated with mRNA splicing and processing (e.g., *Rbm3/25, Rbmx, Cirbp, Srsf5, Luc7l2, Tia1, Snrnp70*, *Fus*) were markedly downregulated by sleep deprivation. In the context of memory impairment by SD, this pattern suggests that although CA1 neurons may initiate synaptic remodeling during SD, disruptions in RNA processing and translational control could ultimately impair effective memory consolidation.

Beyond previously reported DEGs identified from excitatory neurons from the entire hippocampus (31), we further detected (Fig. 6) increased translation of genes involved in insulin signaling and insulin resistance pathways, including *Akt2, Gsk3b, Irs1/2, Pik3r3, Ppp1r3c, Prkab2, Ptpn1/11, Ptprf, and Rps6ka2*. Brain insulin signaling regulates multiple cellular processes including glucose metabolism, cell survival, and synaptic plasticity through PI3K-Akt signaling and downstream phosphorylation of targets such as glycogen synthase kinase 3 beta (GSK-3β) and mTOR (66). Disruption of hippocampal insulin signaling such as developing insulin resistance impairs neuroplasticity and spatial memory in rats (67) and overactivation of GSK-3β has been linked to inhibition of hippocampal long-term potentiation (68). Consistent with this framework, our fosTRAP-seq data indicate that insulin response pathways, particularly those relate to insulin resistance, are substantially altered in SD-sensitive CA1 neurons. These disruptions may serve as a central mechanism linking SD to interrupted synaptic plasticity, impaired signal transmission and reduced memory performance. Importantly, GSK-3β also participates in different aspects and pathways relevant to the onset and development of Alzheimer’s Disease neuropathology via directly promoting tau hyper-phosphorylation, amyloid production and accumulation, and inflammatory molecules (69). Additionally, we observed increased translation of *Ttbk1/2* (tau tubulin kinase 1 and 2), kinases that phosphorylate tau at multiple sites and contribute to tauopathy in neurodegenerative disorders (70). Given that chronic sleep restriction causes hippocampus-dependent memory deficits (71) and promotes tau hyperphosphorylation in both human and rodents (18, 21), our findings suggest a mechanistic continuum whereby repeated acute SD initiates insulin signaling dysfunction and upregulation of tau-related kinases (GSK-3 β, TTBK1/2), which, when sustained, may accelerate tau pathology and neurodegeneration in CA1. Collectively, these results highlight insulin signaling dysfunction as a key molecular feature of SD-induced hippocampal impairment and propose a trajectory linking acute sleep loss to chronic sleep restriction-associated tauopathy and increased vulnerability to neurodegenerative disease.

For future studies, a direct comparison of gene expression profiles between CA1 pyramidal neurons that are sleep loss sensitive (c-Fos+) and insensitive (c-Fos-) during SD will be important for delineating molecular features that confer selective vulnerability to sleep loss. In parallel, the fosTRAP system will enable the manipulation of these SD-sensitive neurons for causal tests of their functional significance. Specifically, determining whether selective activation of this neuronal population is sufficient to induce memory impairments, or whether their inhibition can mitigate SD-induced deficits, will provide critical insight into the circuit mechanisms underlying sleep loss-associated cognitive dysfunction.

In conclusion, this study reveals the subregional impact of sleep deprivation on hippocampal neuronal activation, which highlights the importance of spatially segregated analysis to understand the response to sleep loss. The reactivation of CA1 pyramidal neurons after repeated SD supports the premise that certain excitatory neuron populations are particularly susceptible to repeated sleep loss. Using the c-Fos-RiboTag system, we identified activity-driven gene expression alteration at a high resolution in SD-sensitive CA1 neurons and expanded the understanding of how SD could impair memory consolidation and progress toward chronic sleep restriction induced damages. Our research provides a detailed insight into hippocampal neurons responsive to SD and establishes a foundation for future molecular and functional investigations of neuron subsets activated by sleep loss. Identifying these subsets will help inform strategies to mitigate cognitive deficits associated with SD.

## List of Abbreviation

SD: sleep deprivation
NSD: non-sleep deprivation
CSR: chronic sleep restriction
CA1: Cornus Ammonis 1
CA3: Cornus Ammonis 3
DG: Dentate Gyrus
IEG: immediate early gene
ZT: zeitgeber time
RT: room temperature
h: hour
min: minute
AAV: adeno-associated virus
4-OHT: 4-hydroxytamoxifen
i.p.: intraperitoneally
IHC: immunohistochemistry
PBS: phosphate buffer saline
NP40: nonidet P-40
TRAP: target recombination in active populations
RiboTag: ribosome tagging strategy
TRAPseq: translating ribosome affinity purification sequencing
ANOVA: analysis of variance
DEG: differentially expressed gene
FDR: false discovery rate
GO-BP: gene ontology – biological process
KEGG: Kyoto encyclopedia of genes and genomes
SEM: standard error of the mean

## DECLARATIONS

### Ethics approval and consent to participate

All experiments were conducted in accordance with the standards established by the US National Institutes of Health Guidelines for Animal Care and use was approved by the Institutional Animal Care and Use Committee (IACUC) at the University of Iowa.

### Consent for publication

Not Applicable

### Availability of data and materials

The datasets generated and/or analyzed during the current study are available in the NCBI’s Gene Expression Omnibus repository, GEO Series accession GSE316227, https://www.ncbi.nlm.nih.gov/geo/query/acc.cgi?acc=GSE316227, and DEG accession GSE156925, https://www.ncbi.nlm.nih.gov/geo/query/acc.cgi?acc=GSE156925.

Any additional information underlying this article will be shared on reasonable request to the corresponding author.

### Competing Interests

The authors declare that they have no competing interest.

### Funding

This work was supported by grant funding from the National Institutes on Aging 5R01AG062398 to T.A. and L.C.L. T. A. is supported by a Roy J. Carver Directorship.

### Authors’ contribution

Y.W. and T.A. designed the study with input from E.N.W., J.R., and L.C.L.. Y.W. and E.N.W. performed sleep deprivation and tissue collection. Y.W. performed protein expression experiments and data analysis. Y.W. performed mRNA extraction for fosTRAP-seq and the sequencing data analysis by J.K., C.R., and Y.W.. Y.W. wrote the manuscript with input from all other authors.

## Acknowledgements

The sequencing was performed by Genomics Division in the Iowa Institute of Human Genetics which is supported, in part, by the University of Iowa Carver College of Medicine. We thank Dr. Yong-Seok Lee for his guidance on the CA1 microdissection technique. We also thank Dr. Satya Tadinada and Pravda Quinones-Labernik for their valuable feedback on this manuscript.

## ADDITIONAL FILES

**Additional file1.pdf:**

**Figure S1**. Supplement data for RiboTag expression following single and repeated SD. **A**) Schematic of the c-Fos promoted RiboTag expression following 5h of SD. **B**) Representative image of RiboTag (HAtag) expression in NSD/SD hippocampus. **C**) Comparison of area normalized HAtag+ expression between NSD (n=3) and SD (n=4) groups within area CA1. Unpaired t-test, NSD=48.28±6.60, SD=77.92±19.24, t(6)=0.7073, p>0.05. **D**) Timeline of the CA1 neuron labeling using the cFos-RiboTag strategy (whole hippocampus infusion) with repeated sleep deprivation. **E**) Representative images of neuron labeling shown as RiboTag (HAtag) expression in the NSD and SD hippocampus injected with 15mg/kg or 50mg/kg 4-OHT. **F**) Comparison of neuronal labeling efficacy in hippocampal CA1: area normalized HAtag+ cells in NSD and SD groups injected with 15mg/kg and 50mg/kg of 4-OHT, n= 4-7 mice in each group. Two-way ANOVA test performed. Data shown as mean ± SEM. D: No significant effect of repeated sleep loss (F(1,18)=3.815, p=0.0665) but a significant effect of 4-OHT dose (F(1,18)=13.21, p=0.0019) and a significant interaction of repeat manipulation and 4-OHT dose (F(1,18)=4.049, p=0.0594). Bonferroni post hoc: 15mg/kg: NSD=1145.54±168.88, SD=1209.69±135.90, t(18)=0.0384, p>0.9999; 50mg/kg: NSD=1420.40±139.43, SD=2092.38±328.09, t(18)=3.106, p=0.0366. 15mg/kg vs 50mg/kg: p(NSD)>0.9999, p(SD)=0.0044. **G**) Representative image of background RiboTag expression (∼200/mm2) in negative control mice injected with vehicle (corn oil) rather than 4-OHT.

**Figure S2**. Proximal CA1 shows greater cFos expression than distal CA1 after SD. **A**) Representative image of DAPI, cFos+ and HAtag+ cells in distal vs proximal CA1 in SD-SD group. **B**) Comparison of area normalized DAPI nuclei (1), cFos+ (2), HAtag+ (3), and double labeled (4) cells between distal vs proximal CA1 in SD-SD mice (n=6). Unpaired t-tests. Data shown as mean ± SEM. B1) dCA1=7827.39±190.20, pCA1=7732.98±227.45, t(10)=0.32, *p*=0.7576. B2) dCA1= 776.04±154.53, pCA1=1243.15±140.26, t(10)=2.24, *p*=0.0491. B3) dCA1=1287.53±65.68, pCA1=1993.05±142.29, t(10)=4.50, *p*=0.0011. B4) dCA1=305.80±78.63, pCA1=452.91±41.19, t(10)=1.66, *p*=0.1285. **C**) Comparison of neuronal reactivation rate: percentage (%) of double labeled cells / HAtag+ cells (1) and double labeled cells / cFos+ cells (2) between distal vs proximal CA1 in SD-SD mice. Unpaired t-tests, data shown as mean ± SEM. C1 (%): dCA1=21.97±3.71, pCA1=23.99±1.46, t(10)=0.51, p=0.6230. C2 (%): dCA1=38.43±5.24, pCA1=39.62±3.73, t(10)=0.18, *p*=0.8572.

**Figure S3. fosTRAP-seq biological function enrichment analysis. A, B**) Map of KEGG pathways associated with upregulated DEGs (A) and downregulated DEGs (B) after sleep deprivation in SD-sensitive hippocampal CA1 neurons (adjusted p<0.05). Full details of KEGG pathways see additional file2: Table S4.

**Figure S4. Comparison between fosTRAP-seq and excitatory TRAPseq. A**) Venn diagram representing DEGs specific to fosTRAP-seq analysis. **B**) KEGG pathways (adjusted p<0.05) associated with fosTRAP-seq unique DEGs. **C**) Venn diagram representing DEGs specific to excitatory TRAP-seq analysis. **D**) KEGG pathways (adjusted p<0.05) associated with excitatory TRAPseq unique DEGs. Full details see additional file2: Table S7

**Additional file2.xlsx:**

**Table S1.** Supplement statistical results of 2-way ANOVA analyses of c-Fos expression after single and repeated SD. **Table S2.** List of DEGs revealed by fosTRAP-seq (FDR<0.05, |FC|>1.2). **Table S3.** Enriched GO-BP processes associated with fosTRAPseq DEGs (common pathways enriched in both clusterProfiler and ClueGO analysis). **Table S4.** KEGG biological functions associated with fosTRAP-seq DEGs. **Table S5**. List of common DEGs between fosTRAP-seq and excitatory TRAPseq. **Table S6.** Enriched GO-BP processes associated with common DEGs shared by fosTRAP-seq and excitatory TRAPseq. **Table S7**. Enriched KEGG biological functions of fosTRAP-seq and excitatory TRAPseq unique DEGs.

## Notes

### Competing Interest Statement

The authors have declared no competing interest.

### Summary of Updates

This version of the manuscript has been revised to update the following: Author list updated; Figure 2 is revised to include c-Fos and GAD67 double labeling; the figure of proximal vs distal CA1 activity comparison is moved to supplement material; fosTRAP-seq experiment and gene expression analyses are added as Figure 6, supplement figure 3-4, and supplement table 2-7.

