## Supplementary figures and images for "Repeated sleep deprivation selectively reactivates hippocampal CA1 pyramidal neuron"

### Supplement Figures

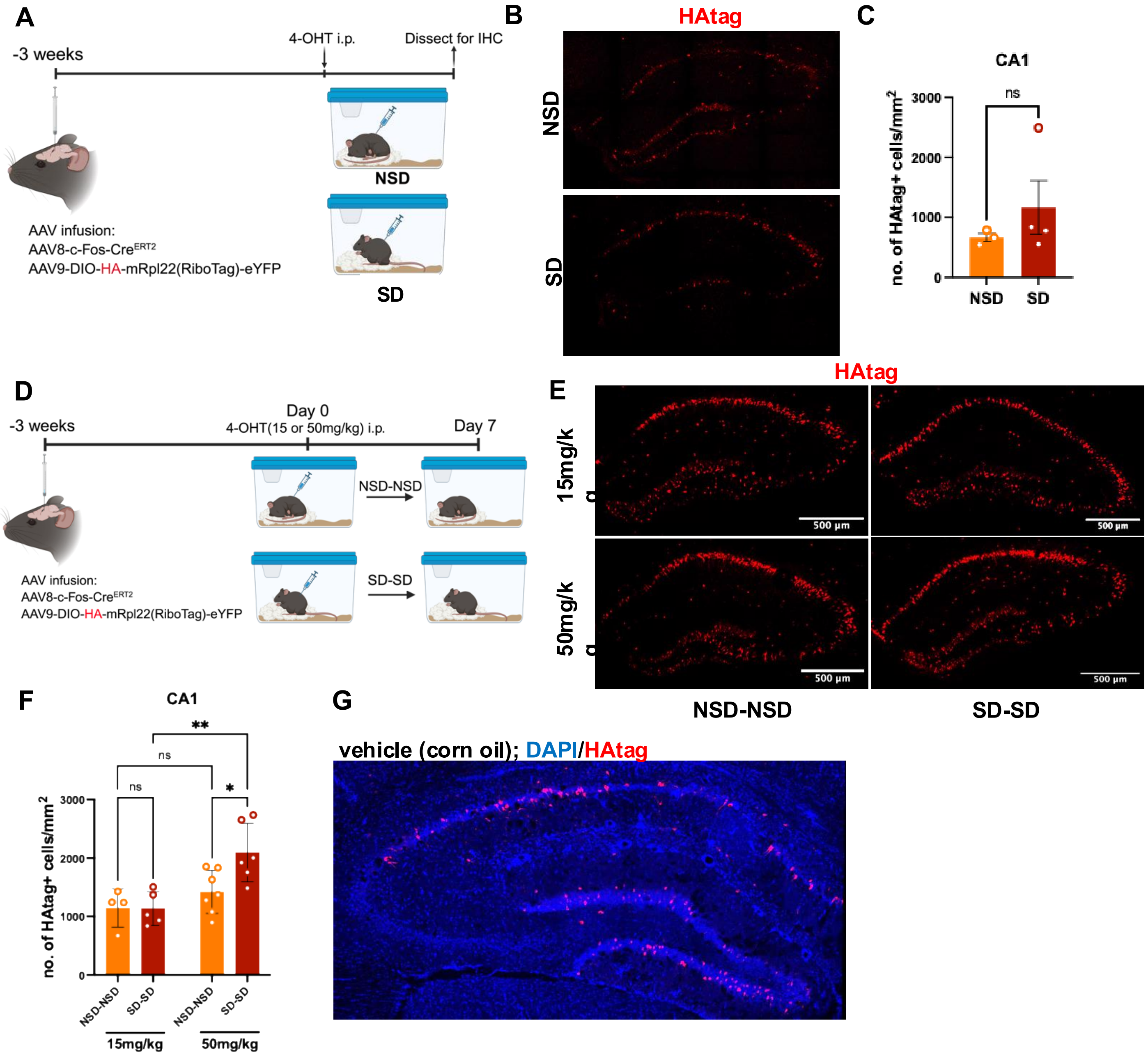

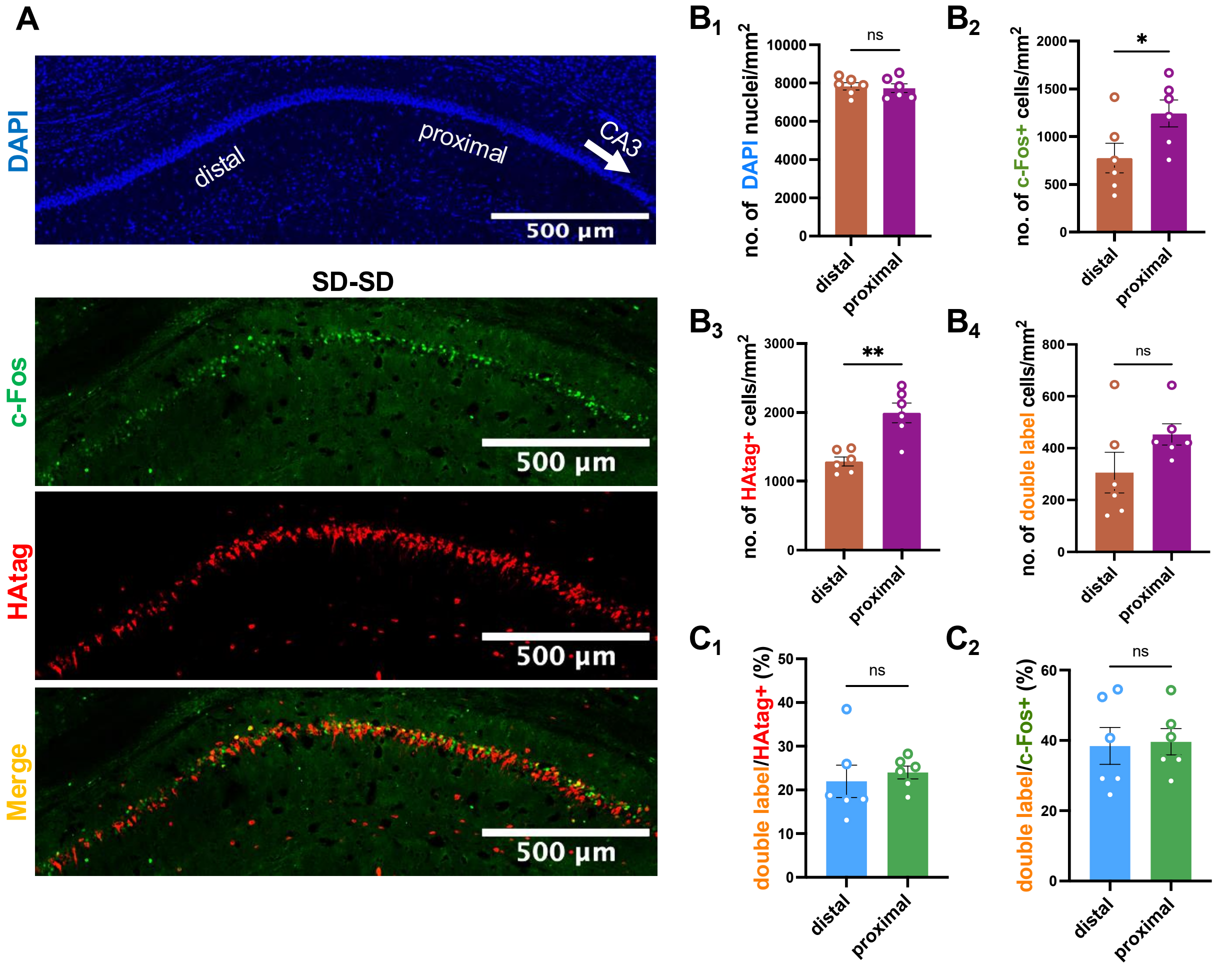

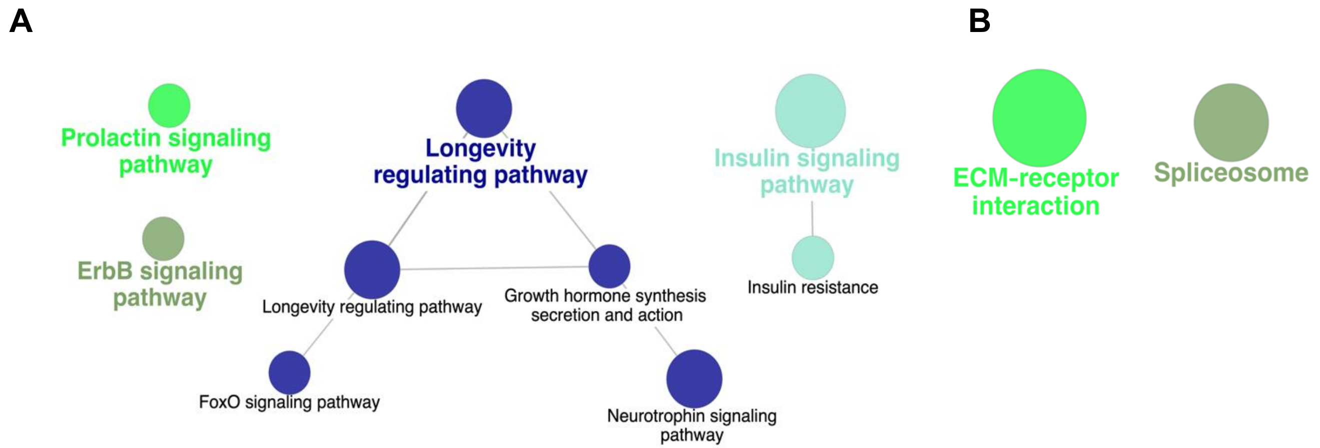

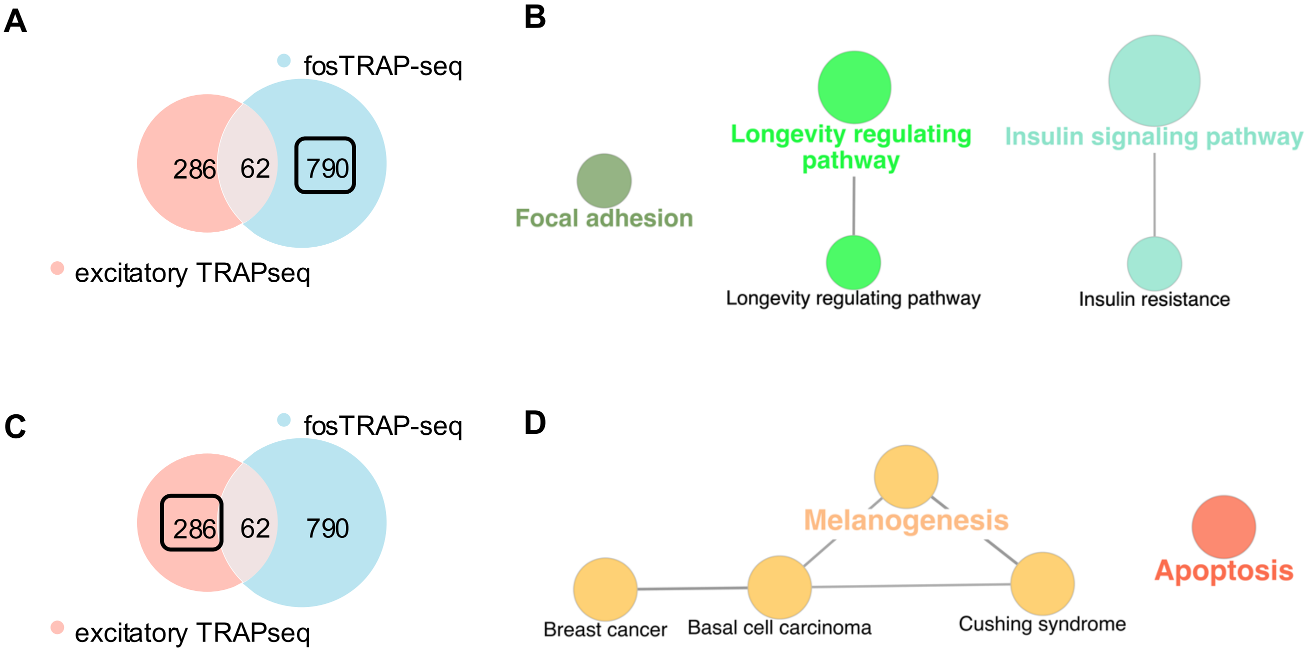
